# Activation of meiotic recombination by nuclear import of the DNA break hotspot-determining complex

**DOI:** 10.1101/2020.06.22.165746

**Authors:** Mélody Wintrebert, Mai-Chi Nguyen, Gerald R. Smith

## Abstract

Meiotic recombination forms crossovers important for proper chromosome segregation and viability of offspring. This process involves many proteins acting at each of the multiple steps of recombination. Recombination is initiated by formation of DNA double-strand breaks (DSBs), which in the several species examined often occur with high frequency at special sites (DSB hotspots). In the fission yeast *Schizosaccharomyces pombe* DSB hotspots are bound with high specificity and activated by linear element (LinE) proteins Rec25, Rec27, and Mug20 which form co-localized nuclear foci with Rec10, essential for all DSB formation and recombination. Here, we identify Rec10’s nuclear localization signal (NLS) and show it is important for coordinated nuclear entry after complex-formation with other LinE proteins. In NLS mutants, recombination is much reduced but not eliminated; nuclear entry of limited amounts of Rec10 can account for the residual recombination. LinEs are related to synaptonemal complex proteins of other species, suggesting that they also may share an as-yet-unidentified NLS and protein complex-formation before nuclear entry.

## Introduction

In sexually reproducing species, one generation gives rise to the next by formation of haploid gametes from diploid somatic cells; fusion of two gametes forms a diploid of the next generation. Gamete formation (meiosis) requires the accurate segregation of each pair of nearly identical chromosomes, one received from each of the previous parents. These chromosomes (homologs) replicate to form sister chromatids, align with each other, and exchange parts to form a physical connection (a crossover) holding the two homologs together. Along with sister chromatid cohesion, crossovers allow tension to form when the homologs begin to segregate from each other, as required for formation of viable gametes. In the absence of crossovers, and thus tension, homologs move nearly randomly in most species tested and often produce gametes with an improper (aneuploid) set of chromosomes which give rise to inviable or disabled progeny.

Formation of crossovers occurs by homologous recombination, which also generates novel combinations of genetic alleles to aid evolution of the species. During meiosis, recombination in the several species examined is initiated by the formation of DNA double-strand breaks [DSBs; reviewed in Nambiar et al. (2019)]; repair of a DSB with an intact homolog can generate a crossover, *i.e.*, reciprocal exchange of homologous chromosome sections flanking the DSB site. In the few species examined DSBs do not occur at random across the genome; rather, they occur at higher than genome-average frequency at special sites, called DSB hotspots (Nambiar et al., 2019). Hotspots are determined by a complex interplay of DNA sequence, histone modification, and binding of special proteins (Keeney et al., 2014). In the fission yeast *Schizosaccharomyces pombe,* the linear element (LinE) proteins Rec25, Rec27, and Mug20 bind DSB hotspots with high specificity and are required for formation of most DSBs at most hotspots (Fowler et al., 2013). These hotspot determinants act with another LinE protein Rec10, which is required for essentially all DSB formation and recombination, *i.e.,* both at hotspots and in the intervening DSB cold regions (Lin and Smith, 1995; Ellermeier and Smith, 2005; Fowler et al., 2013). In the cases tested, these four proteins co-localize in the nucleus and form microscopically visible dots (foci); deletion of any one gene eliminates foci of the others, indicating that the four LinE proteins form a complex in the nucleus (Davis et al., 2008; Estreicher et al., 2012; Fowler et al., 2013). Other analyses, such as co-immunoprecipitation and yeast two-hybrid assays, also indicate that the LinE proteins form a complex (Estreicher et al., 2012).

Previous studies have shown special features of the LinE proteins. Three of them, Rec25, Rec27, and Mug20, contain only 125 – 151 amino acids, whereas Rec10 contains 791 amino acids (Lin and Smith, 1995; Martin-Castellanos et al., 2005; Estreicher et al., 2012). LinE proteins are related to the synaptonemal complex (SC) proteins of other species. Over limited regions, Rec10 shares amino acid sequence similarity with the SC protein Red1 of the budding yeast *Saccharomyces cerevisiae* (Lorenz et al., 2004). Rec27 and Mug20 share amino acid sequence similarity with the SC proteins SYP-2 and DDL-1, respectively, of the round worm *Caenorhabditis elegans* (Fowler et al., 2013). SC proteins hold homologs in alignment from one end to the other, whereas LinE proteins appear by electron microscopy of nuclear spreads to align chromosomes over shorter distances, forming “linear elements” (Bähler et al., 1991). In live cells, however, *S. pombe* meiotic chromosomes also appear aligned side-by-side from one end to the other as seen by light microscopy of the sister chromatid cohesin Rec8 (Ding et al., 2016). The *S. pombe* end-to-end alignment may be more fragile than the SC-mediated alignment of other species and partially fall apart upon opening of cells and spreading the nuclear contents. The recombination-deficiency of both LinE and SC mutants also suggests a similarity in their cellular functions (Page and Hawley, 2004; Cromie and Smith, 2008).

Rec10 has a set of amino acids potentially required for entry of the protein into the nucleus, where it is observed and acts (Lorenz et al., 2004; Spirek et al., 2010; Fowler et al., 2013). This nuclear localization signal (NLS) is predicted to bind to a protein complex involved in nuclear entry of many substrate proteins [reviewed in Soniat and Chook (2015)]. Here, we experimentally identify a two-part NLS in Rec10. Our results show that its alteration or deletion blocks nuclear entry not only of Rec10 but also of other LinE proteins. Our data indicate that the LinEs form a complex even before nuclear entry, and that the other LinE proteins are cargos of Rec10. Without the Rec10 NLS, other LinE proteins form aberrant structures in the cytoplasm. The similarity between *S. pombe* LinEs and the SC of other species suggests that one of the SC proteins may have an NLS, to our knowledge not shown in the SC of any species, and may also form a complex prior to nuclear entry.

## Results

### Identifying the putative NLS of Rec10

Rec10 carries a predicted bipartite nuclear localization signal (NLS) at amino acids 497 to 500 and 516 to 519 (http://nls-mapper.iab.keio.ac.jp/cgi-bin/NLS_Mapper_form.cgi). Other programs, such as PROSITE, CAST, NLStradamus, or NucPred, also predict an NLS in this region. This region is com-posed of clusters of positively charged amino acids, mostly K and R. We designate amino acids 497 to 500 (KRKK) as site A and 516 to 519 (KNKK) as site B (Fig. 1).

**Fig. 1.**
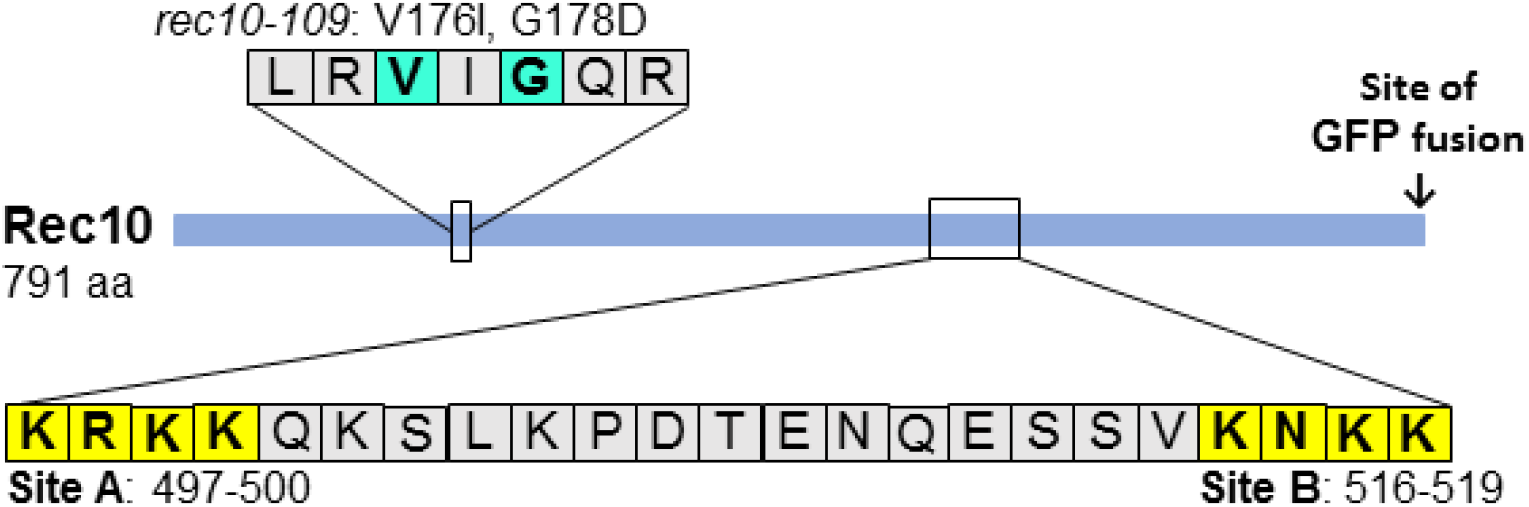
Schematic of Rec10 protein, showing NLS sites A and B and the *rec10-109* mutation discussed here. The solid blue line shows the full-length (791 amino acids) of Rec10. Positions of the NLS sites and their amino acid sequences are shown below, and those of *rec10-109* are shown above.

To test the NLS hypothesis noted above, we mutated site A or B or both, either by deletion or by change of each amino acid to alanine. Alanine is a small, non-charged amino acid, unlike K and R. If these sites are part of the NLS, importin α, part of the nuclear entry complex (Soniat and Chook, 2015), is predicted not to recognize the mutated Rec10 and not to carry it into the nucleus. Mutations were first made in *rec10* on a plasmid and tested for localization of another LinE protein, Rec25, fused to green fluorescent protein (Rec25-GFP) and for recombination proficiency. The NLS mutations were then moved to the chromosome, where *rec10* was fused to the GFP gene to visualize localization of Rec10 itself and to analyze recombination proficiency. Derivatives with chromosomal *rec10* without the GFP fusion were also tested for localization of each of the other LinEs fused to GFP. Similar results were obtained for each *rec10* derivative, indicating that the plasmid *vs* chromosomal location of *rec10* or fusion of Rec10 to GFP does not significantly interact with the NLS.

### Single and double Rec10 NLS mutants have aberrant cytoplasmic localization of linear element proteins

To assess the importance of each putative NLS site in Rec10, we determined the cellular localization of Rec25-GFP, a partner protein of Rec10 (Davis et al., 2008). We tested strains carrying *rec10* mutated (or not) on a plasmid with an estimated copy-number of 1 – 2 (Wahls and Davidson, 2008) (Fig. 2A). For the NLS site A and site B mutants, sharp foci (“speckles”) of Rec25-GFP were prominent in the cytoplasm, and occasional faint dots were in the nucleus. Wild-type Rec10 produced strong nuclear foci of Rec25-GFP without detectable cytoplasmic foci; *rec10Δ* produced only faint background fluorescence, comparable to that in cells without a GFP fusion. Rec25-GFP was readily visible in all other situations (wt and NLS *rec10* mutants), implying that Rec25-GFP is degraded in the absence of Rec10 and likely forms a complex with wt and mutant forms of Rec10 (see Discussion). Similar results were seen with both deletion and Ala mutations at each site and with the double mutations. Rec27-GFP and Mug20-GFP behaved much like Rec25-GFP in cells with wt Rec10 or Rec10 deleted for both NLSs (Fig. 2B and 2C).

**Fig. 2.**
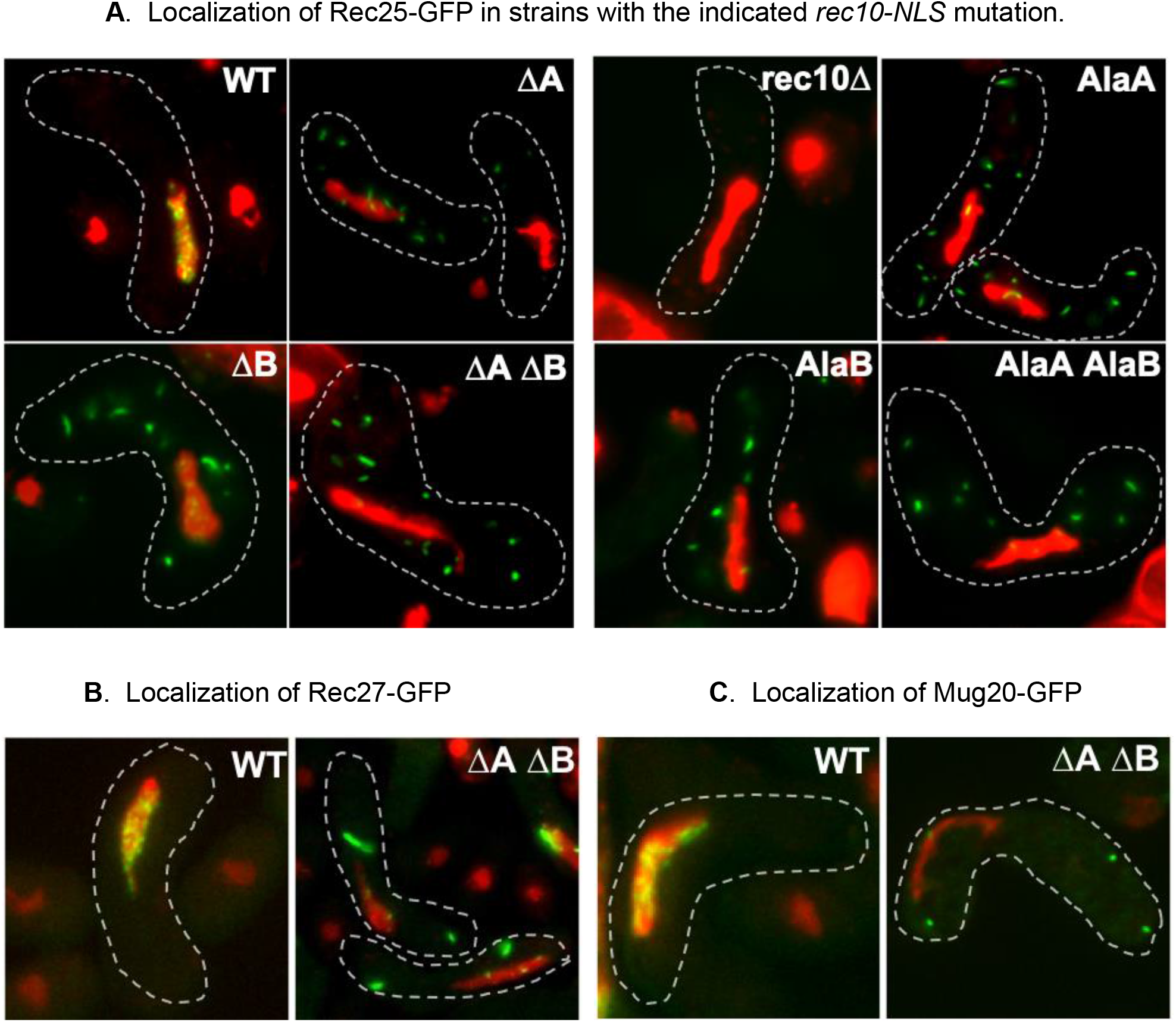
Plasmid-borne *rec10* NLS mutations block entry of other LinE proteins into the nucleus. **(A)** Strains bearing *rec25-GFP* and a complete *rec10* deletion on the chromosome and carrying a plasmid with *rec10^+^* (WT), with the indicated *rec10* NLS mutation, or without *rec10* (rec10Δ) were harvested during asynchronous (*h^90^*) meiosis and examined by light microscopy. Green indicates Rec25, and red indicates chromatin in the nucleus; yellow is their overlap. The dashed line outlines the cell. ΔA indicates deletion of Rec10 NLS site A, and AlaA indicates substitution of alanine for each of the four amino acids of site A; site B was similarly mutated and annotated. Combinations of these mutations are indicated by ΔA ΔB or AlaA AlaB. **(B)** Strains bearing *rec27-GFP* were analyzed as in panel A. **(C)** Strains bearing *mug20-GFP* were analyzed as in panel A.

Based on the cytoplasmic localizations of these three LinE-GFPs, Rec10 appears to aggregate in the cytoplasm when its NLS is mutated (as directly seen below with Rec10-GFP). Rec10 may self-interact (Ma et al., 2017). The cytoplasmic speckles appear to be formed by complexes of Rec10 and the other LinEs, just as they co-localize in the nuclear foci during meiosis (Davis et al., 2008; Estreicher et al., 2012; Fowler et al., 2013). The cytoplasmic speckles in the mutants were less numerous and larger than the nuclear foci in the wild type, suggesting different structures for the cytoplasmic and nuclear LinE complexes (see Discussion).

### Rec10 NLS mutations reduce meiotic recombination

Rec10 is essential for recombination throughout the genome (Ellermeier and Smith, 2005). To test if the NLS, and thus nuclear import, of Rec10 is required for recombination, we measured the frequency of intragenic recombination (gene conversion, or non-reciprocal recombination) at *ade6* and the frequency of intergenic recombination (crossing-over, or reciprocal recombination) between *ade6* and *arg1*. In assays with plasmid-borne *rec10*, conversion was reduced in all the NLS mutants by factors ranging from ~2 to ~6, but in no case did recombination approach the very low level of *rec10Δ* (~300 times lower than that of wild type) (Table 1). Crossovers were reduced by factors of up to 15, but again not as much as in *rec10Δ* (Ellermeier and Smith, 2005). NLS deletions and Ala substitutions behaved similarly. In all four cases, site A mutations reduced recombination more than site B mutations. These results show that the Rec10 NLS is important for recombination but not as essential as Rec10 itself. The residual recombination in *rec10*-NLS mutants may reflect nuclear entry of small amounts of Rec10 (see Discussion).

**Table 1.** *rec10*-NLS mutations reduce but do not eliminate meiotic recombination

| *rec10* genotype | Plasmid NLS | Chromosomal NLS-GFP |
| --- | --- | --- |
| *ade6-M26 x ade6-52* (Ade<sup>+</sup>/10<sup>6</sup> viable spores) <sup>a</sup> |  |  |
| *rec10*<sup>+</sup> | 2900 ± 400 | 1060, 1050 <sup>b</sup> |
| *rec10Δ* | <10 | 3 |
| NLS ΔA | 830 ± 45 | 75, 88 |
| NLS ΔB | 1900 ± 270 | 170, 230 |
| NLS ΔA ΔB | 350 ± 78 | 130, 250 |
| NLS AlaA | 460 ± 75 | 80, 68 |
| NLS AlaB | 1900 ± 380 | 450, 150 |
| NLS AlaA AlaB | 460 ± 46 | 180, 390 |
| *ade6 – arg1* (cM) <sup>c</sup> |  |  |
| *rec10*<sup>+</sup> | 105 | 34 <sup>d</sup> |
| *rec10Δ* | ND <sup>e</sup> | [0/66] <sup>f</sup> |
| NLS ΔA | 24 | 10 |
| NLS ΔB | 60 | 15 |
| NLS ΔA ΔB | 6 | 10 |
| NLS AlaA | 27 | 4 |
| NLS AlaB | 53 | 8 |
| NLS AlaA AlaB | 7 | 6 |
| *ade6 – ura4* (cM) <sup>c</sup> |  |  |
| *rec10*<sup>+</sup> | ND | 90 <sup>g</sup> |
| *rec10Δ* | ND | [0/66] <sup>h</sup> |
| NLS ΔA | ND | 9 |
| NLS ΔB | ND | 40 |
| NLS ΔA ΔB | ND | 19 |
| NLS AlaA | ND | 29 |
| NLS AlaB | ND | 24 |
| NLS AlaA AlaB | ND | 13 |

<sup>a</sup> Data are the mean ± SEM from 3 – 6 crosses, or individual data from 1 or 2 crosses.
<sup>b</sup> 2700 and 2600 in *rec10*<sup>+</sup> without GFP.
<sup>c</sup> Data are from analysis of 120 – 367 (generally 132 – 172) spore colonies, except for individual values (recombinants/spore colonies tested) in square brackets.
<sup>d</sup> 96 cM in *rec10*<sup>+</sup> without GFP.
<sup>e</sup> ND, not determined.
<sup>f</sup> <0.3 cM (0/817) in Ellermeier and Smith (2005).
<sup>g</sup> ~200 in *rec10*<sup>+</sup> without GFP, based on the genome mean of 0.16 cM/kb (Young et al., 2002).
<sup>h</sup> 0.5 cM (2/469) in Ellermeier and Smith (2005).

### Chromosomal Rec10 NLS mutations reflect plasmid-borne mutations in recombination phenotype

To ensure more nearly wild-type levels of Rec10 protein and to assay Rec10 localization itself, we moved the *rec10*-NLS mutations to the chromosome and fused the *rec10* gene to that for GFP (Rec10-NLS-GFP). The NLS mutant proteins formed cytoplasmic speckles, much like those of Rec25-GFP described above for plasmid-borne *rec10* NLS mutations, or larger clumps (Fig. 3). *rec10* NLS deletion and Ala substitution mutations had similar phenotypes, as did the single and double site mutations. The similar appearance of Rec25-GFP (Fig. 2) and Rec10-GFP (Fig. 3) is consistent with these proteins forming a complex, either in the nucleus (wild type) or in the cytoplasm (NLS mutants) (see Discussion).

**Fig. 3.**
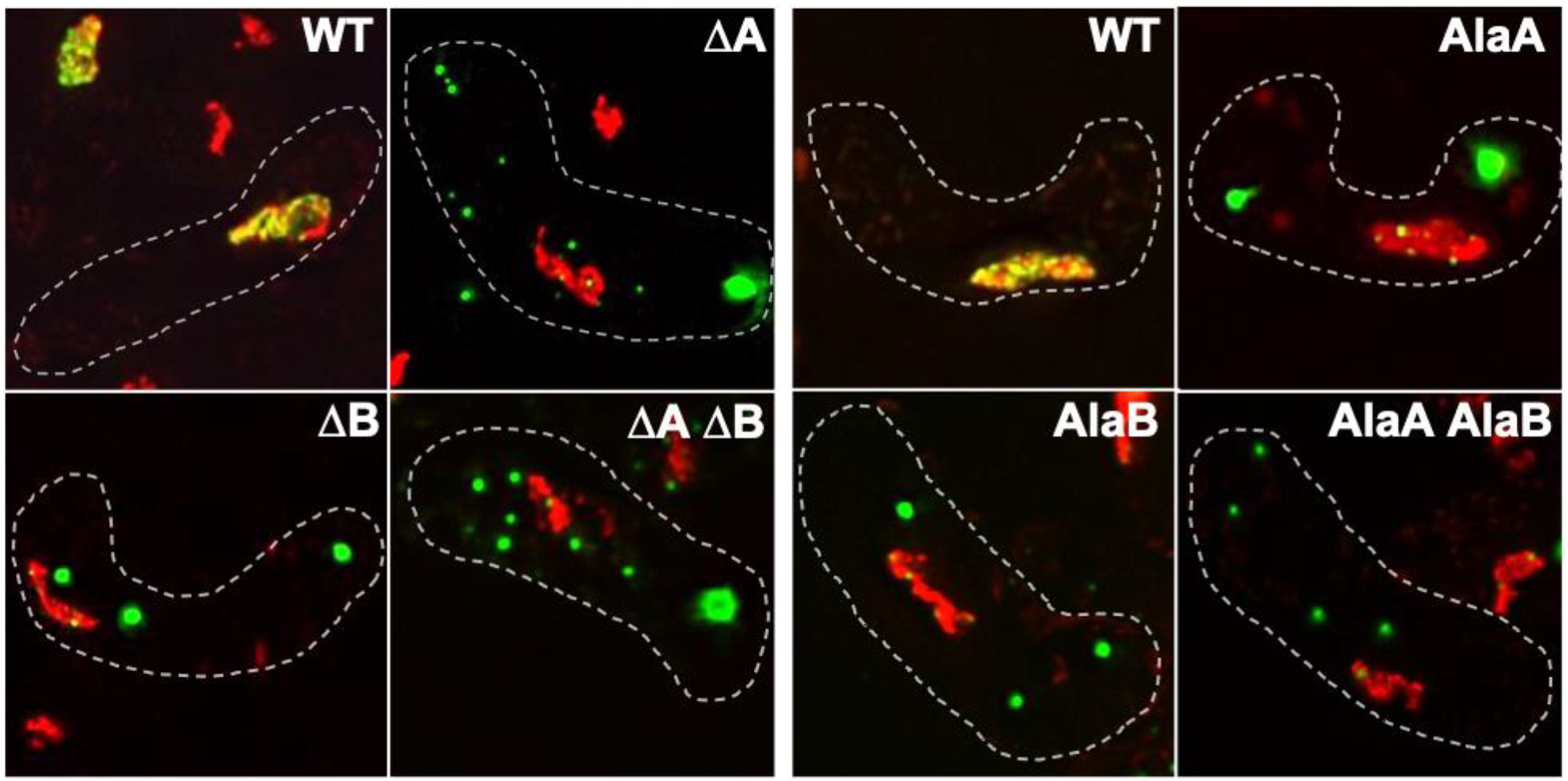
Chromosomal *rec10* NLS mutations block entry of Rec10-GFP into the nucleus. Strains without (WT) or with a *rec10* NLS mutation were harvested during asynchronous (*h^90^*) meiosis and examined by light microscopy. Green indicates Rec10, and red indicates chromatin in the nucleus; yellow is their overlap. The dashed line outlines the cell. NLS mutations are designated as in Fig. 2.

Recombination was also reduced similarly in chromosomal and plasmid-borne *rec10* NLS mutants (Table 1). Recombinant frequencies for *ade6* gene conversion and *ade6 – arg1* crossing-over, as well as for *ade6 – ura4* crossing over, were reduced by factors of ~3 – 8 by the chromosomal NLS mutations, comparable to the reductions by the plasmid-borne alleles. In nine of ten cases tested, site A mutations reduced recombination more than site B mutations; double site A site B mutations reduced recombination at least as much as a single mutation. Recombinant frequencies were ~3 times lower with the chromosomal *rec10* alleles (both wild-type and NLS mutant), in which Rec10 is fused to GFP (Table 1). This fusion, however, has nearly wild-type levels of crossing-over at *lys3 – ura1 – met5* on chromosome 1 (Fowler et al., 2013), suggestive of the regional specificity of certain *rec10* missense mutations (see Discussion).

To compare directly plasmid-borne and chromosomal *rec10*-NLS mutations (without GFP fusion), we assayed Rec25-GFP localization in both sets of strains. For each single and double mutant, Rec25-GFP formed abundant cytoplasmic speckles and was depleted in the nucleus (Figs. 2 and 4). As with the plasmid-borne mutations, deletion and Ala substitution mutations, as well as single and double mutations, behaved similarly. This outcome is as expected if the *rec10* gene is regulated similarly on the low copy-number plasmid and on the chromosome.

**Fig. 4.**
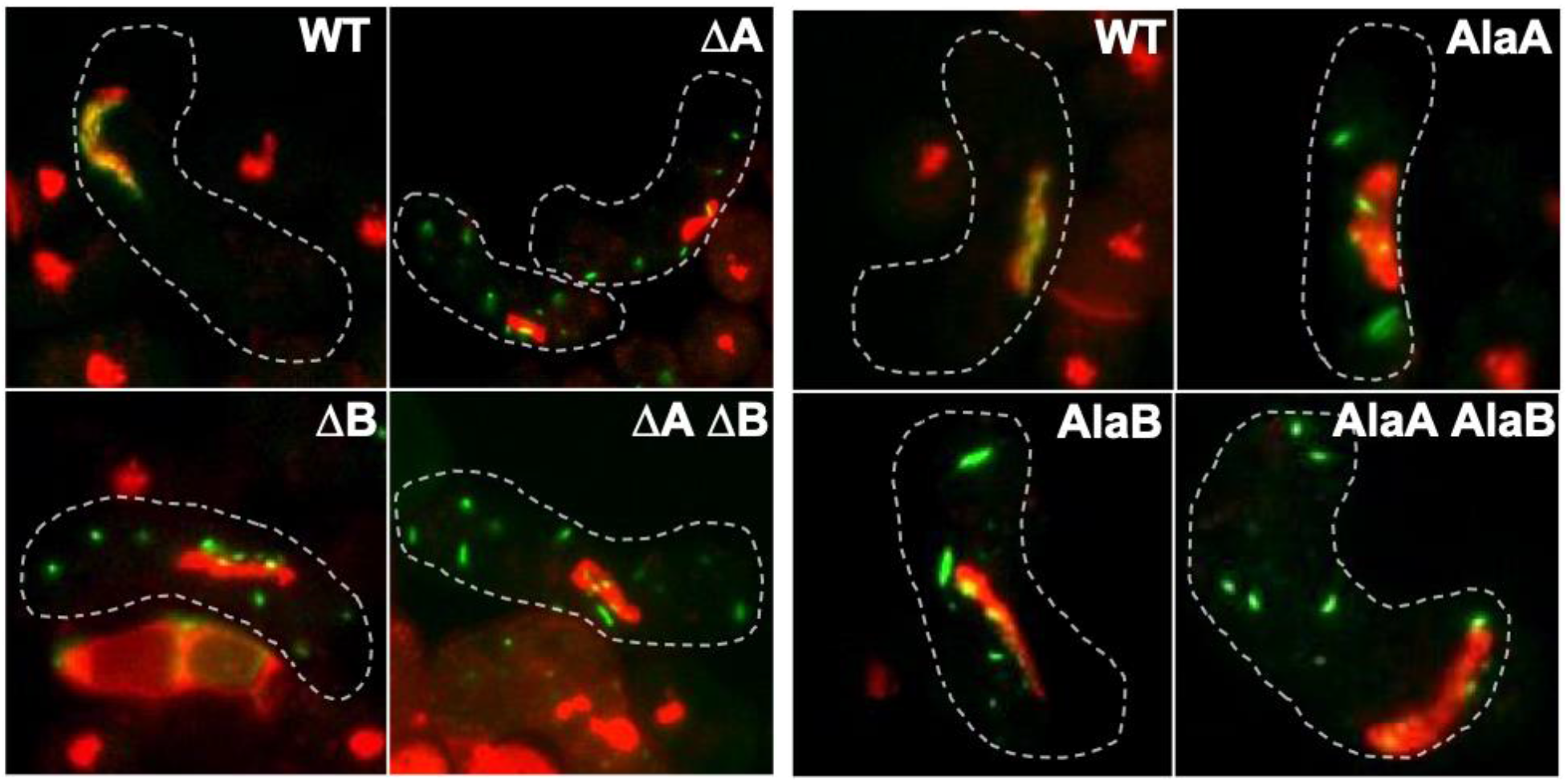
Chromosomal *rec10* NLS mutations block entry of Rec25-GFP into the nucleus. Strains bearing *rec25-GFP* and without (WT) or with a *rec10* NLS mutation were harvested during asynchronous (*h^90^*) meiosis and examined by light microscopy. Green indicates Rec25, and red indicates chromatin in the nucleus; yellow is their overlap. The dashed line outlines the cell. NLS mutations are designated as in Fig. 2.

### Rec10 abundance is not altered by NLS mutations

Retention of Rec10 in the cytoplasm rather than entry into the nucleus could lead to its partial degradation and reduce, but not eliminate, meiotic recombination. To test this possibility, we induced into meiosis strains carrying the chromosomal *rec10* alleles with the NLS mutations and the GFP fusion. Protein abundance was assayed by polyacrylamide gel electrophoresis and Western blot analysis using antibody to GFP. The results (Fig. 5) show that Rec10 is meiotically induced to the same level in wild-type strains and in each of the six NLS mutants. Thus, the relocalization of Rec10-GFP and Rec25-GFP and the reduction of recombination cannot be attributed to aberrant expression of *rec10* or instability of Rec10-GFP protein

**Fig. 5.**
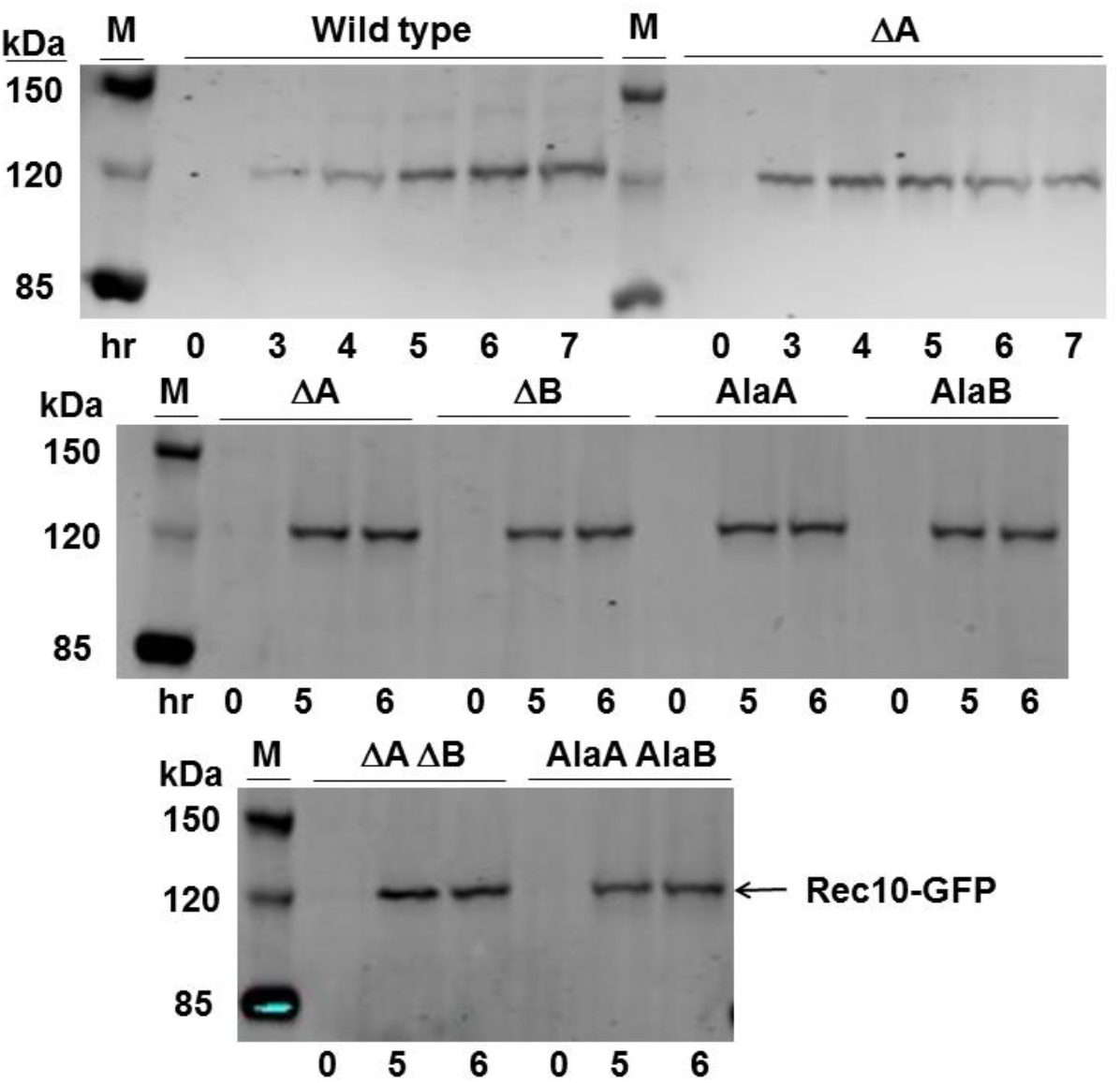
Rec10 protein is present at wild-type levels in *rec10-NLS* mutants. Strains with chromosomal *rec10-NLS-GFP* fusions were induced for meiosis (*pat1-as1*) and analyzed by Western blots for Rec10 abundance in cell extracts. NLS mutations are designated as in Fig. 2. M, protein markers with the indicated masses.

## Discussion

Our results show the importance of the nuclear localization signal (NLS) of Rec10 for entry of LinE proteins into the nucleus and their promotion of recombination. These results imply that Rec10 forms a complex with the other LinE proteins before nuclear entry. Based on these and earlier data, we also infer that alterations to Rec10 structure have differential effects on its activity in different chromosomal regions.

Both of the two predicted Rec10 NLS sites are important for Rec10 and other LinE proteins to enter the nucleus. Site A, amino acids KRKK (497 – 500), appears to be more important than site B, amino acids KNKK (516 – 519) (Fig. 1), at least for recombination, the most quantitative assay used here (Table 1). The double mutant, whether deletions or substitutions to alanine, behaved like single site mutants, suggesting that the two sites act together. Two-site NLSs in other proteins behave similarly (Soniat and Chook, 2015), suggesting that Rec10’s NLS is canonical. Rec25 was visible at only low levels in the nucleus in the NLS mutants, showing that its nuclear entry mainly depends on Rec10’s NLS. Rec10 and the other LinEs co-localize in the nucleus in wild-type Rec10 strains (Davis et al., 2008; Estreicher et al., 2012; Fowler et al., 2013). The simplest interpretation of these results is that Rec10 forms a complex with the other LinEs in the cytoplasm and only then does this complex enter the nucleus. In NLS mutants this complex still forms but remains mainly in the cytoplasm, consistent with the similar appearance of Rec10-GFP and Rec25-GFP cytoplasmic speckles in the NLS mutants (compare Figs. 2 and 4 with Fig. 3). These speckles appear qualitatively larger and less numerous than the nuclear foci in wild-type NLS strains. We suppose that the cytoplasmic NLS complex forms aggregates when it cannot enter the nucleus, much as SC proteins of other species form aggregates (“polycomplexes”) when they cannot form proper nuclear structures (Sym and Roeder, 1995). LinE proteins share amino acid sequence similarity with SC proteins (see Introduction), consistent with this view.

Residual recombination in all the NLS mutants implies that a small amount of Rec10 still enters the nucleus in these mutants. The LinE complex, with a MW of 139.0 kDa assuming one of each subunit, may be unable to enter the nucleus without the aid of importin, while Rec10 alone, with a MW of 89.9 kDa, may be able to enter, as shown for other proteins in this size range (Wang and Brattain, 2007). Complete deletion of *rec10* reduces recombination as much as does deletion of the DSB-forming protein Rec12, up to 500 times less recombination than in wild-type cells (DeVeaux et al., 1992; Ellermeier and Smith, 2005). Residual recombination in the Rec10 NLS mutants may occur without participation of the other LinE proteins (Rec25, Rec27, and Mug20), because these proteins are not as stringently required for recombination as Rec10 (Davis et al., 2008; Estreicher et al., 2012). In the NLS mutants, nuclear Rec10 levels may be below or near the level of detection by microscopy but still be sufficient to promote low, but detectable, levels of recombination.

We noted that the Rec10-GFP fusion, compared to wild-type Rec10, reduced recombination on chromosome 3 (*ade6, ade6 – arg1,* and *ade6 – ura4*) (Table 1), although it does not significantly reduce recombination at two adjacent intervals on chromosome 1 (*lys3 – ura1 – met5*) (Fowler et al., 2013). This regional specificity is similar to that of *rec10-109*, a double missense mutation (V176I, G178D) in the N-terminal region of Rec10 (DeVeaux and Smith, 1994; Ma et al., 2017) (Fig. 1). Both mutants retain nearly wild-type levels of recombination in the chromosome 1 intervals tested but have reduced recombination at *ade6* and in the *ade6 – arg1* and *ade6 – ura4* intervals on chromosome 3 (DeVeaux and Smith, 1994; Fowler et al., 2013) (Table 1). The GFP-dependent reductions were nearly the same whether or not Rec10 was mutated at the NLS, indicating that the effect is independent of the amount of Rec10 in the nucleus and whether or not it is part of the LinE complex. Although the basis for this regional specificity has not been determined, it likely reflects the abnormal structure of Rec10, and thus the LinE complex, in these mutants. The altered LinE structure may bind to some DSB hotspots (*e.g.,* on chromosome 1) but not to others (*e.g.,* on chromosome 3).

The bipartite NLS determined here is conserved among *Schizosaccharomyces* and other species. Both parts of the NLS are present, with only small differences, at the same positions in an alignment of *S. kambucha, S. octosporus* and *S. cryophilus* (Ma et al., 2017). The more distantly related *S. japonicus* has a bipartite NLS, predicted by NLS Mapper, about 70 amino acids toward the N-terminus. This outcome suggests that these species, which contain orthologs of all four LinE proteins, also control DSB hotspot activity by formation and nuclear import of a LinE protein complex. Although *S. cerevisiae* Red1 has a bipartite NLS, predicted by NLS Mapper, at amino acids 551 – 574 (of 827 total), this has not, to our knowledge, been experimentally verified. The fruit fly *Drosophila melanogaster* SC protein C(2)M contains a predicted bipartite NLS (Manheim and McKim, 2003), but this NLS has not been confirmed experimentally (K. McKim, pers. comm). Making and testing NLS-deficient mutants in other species would allow determining the extent to which formation of a large protein complex is required for nuclear entry of proteins critical for meiotic recombination.

## Materials and methods

### Strains, plasmids, and oligonucleotides

*S. pombe* strains and their genotypes are in Table S1; plasmids are in Table S2. Oligonucleotides, purchased from Integrated DNA Technologies and used for mutagenesis and DNA sequencing, are in Table S3. Genetic manipulations and culture media were as described (Smith, 2009),

### Mutagenesis and strain constructions

Q5 Site-Directed Mutagenesis Kit (New England BioLabs) was used to make substitutions, and Quickchange Site-Directed Mutagenesis Kit (Agilent) was used to make deletions, following the protocols provided by the companies. Mutagenesis was done on pYL176, a plasmid carrying *rec10*, an ampicillin-resistance gene for selection in *E. coli,* and the *S. cerevisiae URA3* gene for selection in *S. pombe ura4* mutants (Lin and Smith, 1995). Plasmids were extracted from four to eight separate *E. coli* transformants for each mutant and sequenced. Plasmids with the desired mutations were introduced into *S. pombe* strain GP9806 (*h^90^ rec10-175::kanMX6 ura4-294 rec25-303::GFP-hphMX6*) by transformation to Ura^+^. Table S4 lists the NLS mutations, the oligonucleotides for their construction, the initial plasmids, and the chromosomal derivatives.

The *rec10-NLS* mutations were transferred to the chromosome by digestion of these plasmids with *Mlu*I and *Nsi*I to create a 2.67 kb fragment containing the *rec10* gene and ~0.1 kb to each side. The digested DNA was used to transform strain GP7301 (*rec10-260::ura4^+^*) to 5-fluoro-orotic acid (FOA)-resistance (Boeke et al., 1984).

To create *rec10-NLS-GFP* chromosomal derivatives, strain GP9845 (*rec10-306::ura4^+^-GFP-natMX6*) was similarly transformed to FOA-resistance, using the 2.17 kb *Mlu*I - *Xho*I fragment containing most of *rec10* and its NLS mutation. *rec10-306* was made in two steps – substitution of *natMX6* for *kanMX6* (in *rec10-203::GFP-kanMX6* to make *rec10-301::GFP-natMX6*) (Hentges et al., 2005), followed by substitution of *ura4^+^* for the *rec10* NLS region. For the second substitution, DNA flanking the left and right sides of the Rec10 NLS sites was generated with oligonucleotides OL4278 and OL4279 (left) and OL4280 and OL4281 (right) using *S. pombe rec10^+^* DNA as template; OL4279 and OL4280 have 20 nucleotides of *ura4^+^* sequence. *ura4^+^* DNA was generated by a PCR with OL4282 and OL4283 using pFY20 as template; OL4282 and OL4283 have 20 nucleotides of *rec10^+^* sequence. The three PCR reactions were diluted 1:10 and used as template in a PCR using OL4278 and OL4281, which generated the *rec10* coding sequence substituted from nucleotides 373 to 1632 with ~1.8 kb bearing *ura4^+^*. Strain GP9836 (*rec10-301::GFP-natMX6*) was transformed with this DNA, with selection for Ura^+^ to generate the chromosomal *rec10-306::GFP-natMX6* allele. *rec10-306* has the *GFP* gene fused to the C-terminus of the *rec10* gene and the nourseothricine-resistance determinant *natMX6* inserted near the 3’ end of the *GFP* gene (Fowler et al., 2013). Integration at *rec10* was verified by PCR specific for *rec10^+^* and *rec10-306* (Table S3).

Drug-resistance genes nourseothricin N-acyltransferase (*nat*) and hygromycin B phosphotransferase (*hph*) were PCR-amplified from plasmids pFA6a-*natMX6* and pFA6a-*hphMX6* (Table S2) using oligonucleotides and conditions similar to those in Sato *et al.* (2005). The resultant fragments were used to transform strains to the new drug-resistance, thereby replacing the resident drug-resistance gene (Bähler et al., 1998). Strains GP8762 (*h^90^ rec10-203::GFP-kanMX6*) and GP8766 (*h^90^ rec25-204::GFP-kanMX6*) were transformed with *natMX6* and *hphMX6*, respectively, to make strains GP9747 (*h^90^ rec10-301::GFP-natMX6*) and GP9745 (*h^90^ rec25-303::GFP-hphMX6*).

### Recombination assays

#### Plasmid-borne rec10-NLS mutations

Strain GP6994 (*h^−^ ade6-52 ura4-D18 rec10-175::kanMX6*) and GP9775 (*h^+^ ade6-M26 ura4-D18 rec10-175::kanMX6 arg1-14*) were transformed with the mutated pYL176 plasmids. For each mutant and pYL176 (*rec10*^+^) control, the two strains were crossed. After two days on sporulation medium (SPA) plus adenine and arginine, spores were harvested, treated sequentially with glusulase and ethanol, and washed. Appropriate dilutions of spore suspensions were spread on yeast extract medium (YEA) medium, to assay total viable spore titer, and on YEA medium supplemented with guanine (YEAG), which inhibits adenine uptake, to assay Ade^+^ recombinant titers; Ade^+^ recombinant frequency is expressed as the ratio of these titers. Colonies were picked from YEA, noting their color (light red for *ade6-52* and dark red for *ade6-M26*) to YEA plus adenine medium and replicated with velvet onto minimal nitrogen base agar (NBA) plus adenine medium supplemented or not with arginine to score *arg1*. Intergenic recombinant frequencies were converted to genetic distance using Haldane’s equation. See Smith (2009) for details.

#### Chromosomal rec10-NLS mutations

To test *rec10-GFP* NLS mutants, FOA-resistant transformants of strain GP9845 (*h^90^ rec10-306::ura4^+^-GFP-natMX6 ura4-D18*) were mated with strain GP4914 (*h^+^ ura4-D18 ade6-M26 arg1-14)* to obtain *h^+^ ura4-D18 ade6-M26 arg1-14 rec10-NLS-GFP-natMX6* derivatives. These isolates (Table S4) were mated with strain GP4625 (*h^−^ ade6-52 rec10-175::kanMX6*). Control strains were GP4914 (*rec10*^+−^) and GP9775 (*rec10-175::kanMX6*). Spores were analyzed as above plus replication onto minimal NBA plus adenine and arginine medium supplemented or not with uracil to score *ura4*.

### Fluorescence microscopy

Homothallic (*h^90^*) cells were spotted on un-supplemented SPA; after 16 hr of incubation at 25 °C, cells were harvested, stained with Hoechst 33342, and spotted on poly-lysine coated microscope slides. Cells with an elongated nucleus (in the horsetail stage) were scored for localization of green fluorescence (GFP). Images are of a single focal plane (Figs. 2 and 4) or maximal projections of 9 –12 sections, step size 0.2 μm, to cover the whole cell (Fig. 3). Images were obtained with an Evos FL Auto 2 microscope (ThermoFisher Scientific) at 60 x magnification (Figs. 2 and 4) or DeltaVision microscope (GE Healthcare) at 100 x magnification (Fig. 3). Images taken by DeltaVision microscope were deconvolved and projected using softWorX software program (Applied Precision). All images were analyzed with ImageJ software.

### Protein abundance by Western blot analysis

Synchronous meiosis was induced as described by Hyppa and Smith (2009) using the *pat1-as1* (L95G) allele at 25 °C (Guerra-Moreno et al., 2012). In brief, haploid *h^−^ pat1-as1* (L95G) *rec10-NLS-GFP* strains were grown in liquid minimal medium EMM2, washed, and starved for nitrogen. After 18– 22 hr of starvation, when the cultures had reached OD_600_ of 0.6 – 0.8, NH_4_Cl and 3-MB-PP1 (SigmaAldrich) were added to 0.50 % (w/v) and 25 μM, respectively, to start meiosis. Culture samples (25 mL) were collected at the indicated times. Cells were collected by centrifugation and washed once with water and once with 20% trichloroacetic acid (TCA). Cell pellets were frozen on liquid nitrogen and kept at −20 °C until protein extraction. Proteins were extracted using TCA and glass beads. Protein extracts were suspended in 50 μL of NuPAGE LDS Sample buffer (ThermoFisher Scientific). Samples (5 μL each) were loaded into wells of a NuPAGE Tris-acetate 3 – 8% polyacrylamide gel (ThermoFisher Scientific) and electrophoresed for 60 min at 15 volts/cm. Proteins were transferred to Immobilon-FL PVDF membranes (Millipore), which were probed sequentially with rabbit Clontech Living Colors Full Length Anti-GFP antibodies (Takara Bio) and goat 680 anti-rabbit IgG antibodies (LiCor). Membrane images were taken by Li-Cor Odyssey NIR Scanner.

## Acknowledgments

We are grateful to Emma Chen for excellent technical assistance; Yu-Chien Chuang and Randy Hyppa for construction of strain GP9845 (*rec10-306::ura4^+^-GFP-natMX6*) and much helpful advice; Yu-Chien Chuang for help with microscopy; Kim McKim and Scott Hawley for information about *D. melanogaster* C(2)M and other SC proteins; and Sue Amundsen for helpful comments on the manuscript. This work was supported by research grant R35 GM118120 from the National Institutes of General Medical Sciences from the United States of America and philanthropic contributions to the Fred Hutchinson Cancer Research Center.

## References

Bähler, J., P. Schuchert, C. Grimm, and J. Kohli. 1991. Synchronized meiosis and recombination in fission yeast: observations with *pat1-114* diploid cells. Current Genetics. 19:445–451.

Bähler, J., J.-Q. Wu, M.S. Longtine, N.G. Shah, A. McKenzie III, A.B. Steever, A. Wach, P. Philippsen, and J.R. Pringle. 1998. Heterologous modules for efficient and versatile PCR-based gene targeting in *Schizosaccharomyces pombe*. Yeast. 14:943–951.

Boeke, J.D., F. LaCroute, and G.R. Fink. 1984. A positive selection for mutants lacking orotidine-5’-phosphate decarboxylase activity in yeast: 5-fluoro-orotic acid resistance. Mol Gen Genet. 197:345–346.

Cromie, G.A., and G.R. Smith. 2008. Meiotic recombination in *Schizosaccharomyces pombe*: A paradigm for genetic and molecular analysis. In Recombination and meiosis: Models, means, and evolution. R. Egel and D.-H. Lankenau, editors. Springer-Verlag, Berlin. 195–230.

Davis, L., A.E. Rozalén, S. Moreno, G.R. Smith, and C. Martin-Castellanos. 2008. Rec25 and Rec27, novel components of meiotic linear elements, link cohesin to DNA breakage and recombination in fission yeast. Current Biology. 18:849–854.

DeVeaux, L.C., N.A. Hoagland, and G.R. Smith. 1992. Seventeen complementation groups of mutations decreasing meiotic recombination in *Schizosaccharomyces pombe*. Genetics. 130:251–262.

DeVeaux, L.C., and G.R. Smith. 1994. Region-specific activators of meiotic recombination in *Schizosaccharomyces pombe*. Genes Development. 8:203–210.

Ding, D.Q., A. Matsuda, K. Okamasa, Y. Nagahama, T. Haraguchi, and Y. Hiraoka. 2016. Meiotic cohesin-based chromosome structure is essential for homologous chromosome pairing in *Schizosaccharomyces pombe*. Chromosoma. 125:205–214.

Ellermeier, C., and G.R. Smith. 2005. Cohesins are required for meiotic DNA breakage and recombination in *Schizosaccharomyces pombe*. Proc. Natl. Acad. Sci. USA. 102:10952–10957.

Estreicher, A., A. Lorenz, and J. Loidl. 2012. Mug20, a novel protein associated with linear elements in fission yeast meiosis. Current Genetics. 58:119–127.

Fowler, K.R., S. Gutiérrez-Velasco, C. Martín-Castellanos, and G.R. Smith. 2013. Protein determinants of meiotic DNA break hotspots. Mol. Cell. 49:983–996.

Guerra-Moreno, A., I. Alves-Rodrigues, E. Hidalgo, and J. Ayte. 2012. Chemical genetic induction of meiosis in *Schizosaccharomyces pombe*. Cell Cycle. 11:1621–1625.

Hentges, P., B. Van Driessche, L. Tafforeau, J. Vandenhaute, and A.M. Carr. 2005. Three novel antibiotic marker cassettes for gene disruption and marker switching in *Schizosaccharomyces pombe*. Yeast. 22:1013–1019.

Hyppa, R.W., and G.R. Smith. 2009. Using *Schizosaccharomyces pombe* meiosis to analyze DNA recombination intermediates. In Meiosis.S. Keeney, editor. Humana Press, Totowa, NJ. 235–252.

Keeney, S., J. Lange, and N. Mohibullah. 2014. Self-organization of meiotic recombination initiation: general principles and molecular pathways. Annu Rev Genet. 48:187–214.

Lin, Y., and G.R. Smith. 1995. Molecular cloning of the meiosis-induced *rec10* gene of *Schizosaccharomyces pombe*. Current Genetics. 27:440–446.

Lorenz, A., J.L. Wells, D.W. Pryce, F.E. Novatchkova, F. Eisenhaber, R.J. McFarlane, and J. Loidl. 2004. *S. pombe* meiotic linear elements contain proteins related to synaptonemal complex components. J. Cell Sci. 117:3343–3351.

Ma, L., K.R. Fowler, C. Martin-Castellanos, and G.R. Smith. 2017. Functional organization of protein determinants of meiotic DNA break hotspots. Sci Rep. 7:1393.

Manheim, E.A., and K.S. McKim. 2003. The synaptonemal complex component C(2)M regulates meiotic crossing over in Drosophila. Curr Biol. 13:276–285.

Martin-Castellanos, C., M. Blanco, A.E. Rozalen, L. Perez-Hidalgo, A.I. Garcia, F. Conde, J. Mata, C. Ellermeier, L. Davis, P. San-Segundo, G.R. Smith, and S. Moreno. 2005. A large-scale screen in *S. pombe* identifies seven novel genes required for critical meiotic events. Current Biology. 22:2056–2062.

Nambiar, M., Y.C. Chuang, and G.R. Smith. 2019. Distributing meiotic crossovers for optimal fertility and evolution. DNA Repair (Amst). 81:102648.

Page, S.L., and R.S. Hawley. 2004. The genetics and molecular biology of the synaptonemal complex. Annual Review of Cell and Developmental Biology. 20:525–558.

Sato, M., S. Dhut, and T. Toda. 2005. New drug-resistant cassettes for gene disruption and epitope tagging in *Schizosaccharomyces pombe*. Yeast. 22:583–591.

Smith, G.R. 2009. Genetic analysis of meiotic recombination in *Schizosaccharomyces pombe*. In Meiosis.S. Keeney, editor. Humana Press, Totowa, NJ. 65–76.

Soniat, M., and Y.M. Chook. 2015. Nuclear localization signals for four distinct karyopherin-beta nuclear import systems. Biochem J. 468:353–362.

Spirek, M., A. Estreicher, E. Csaszar, J.L. Wells, R.J. McFarlane, F.Z. Watts, and J. Loidl. 2010. SUMOylation is required for normal development of linear elements and wild-type meiotic recombination in *Schizosaccharomyces pombe*. Chromosoma. 119:59–72.

Sym, M., and G.S. Roeder. 1995. Zip1-induced changes in synaptonemal complex structure and polycomplex assembly. J Cell Biol. 128:455–466.

Wahls, W.P., and M.K. Davidson. 2008. Low-copy episomal vector pFY20 and high-saturation coverage genomic libraries for the fission yeast *Schizosaccharomyces pombe*. Yeast. 25:643–650.

Wang, R., and M.G. Brattain. 2007. The maximal size of protein to diffuse through the nuclear pore is larger than 60kDa. FEBS letters. 581:3164–3170.

Young, J.A., R.W. Schreckhise, W.W. Steiner, and G.R. Smith. 2002. Meiotic recombination remote from prominent DNA break sites in *S. pombe*. Mol. Cell. 9:253–263.

