## Supplemental information for "Activation of meiotic recombination by nuclear import of the DNA break hotspot-determining complex"

**Table S1. *S. pombe* strains**[a]

| Strain number | Genotype[b] | Used for, or to construct strains for |
|---|---|---|
| GP50 | *h*$^{90}$ | Microscopy (background control) |
| GP4625 | *h*$^{-}$ *ade6-52 rec10-175::kanMX6* | Recombination assay, chromosomal *rec10::GFP* |
| GP4914 | *h*$^{+}$ *ura4-D18 ade6-M26 arg1-14* | Recombination assay, chromosomal *rec10::GFP* |
| GP6994 | *h*$^{-}$ *rec10-175::kanMX6 ura4-D18 ade6-52* | Recombination assay, plasmid-borne *rec10* |
| GP9747 | *h*$^{90}$ *rec10-301::GFP-natMX6* | Microscopy, Rec10 localization |
| GP9775 | *h*$^{+}$ *rec10-175::kanMX6 ura4-D18 ade6-M26 arg1-14* | Recombination assay, plasmid-borne *rec10* |
| GP9806 | *h*$^{90}$ *rec10-175::kanMX6 rec25-303::GFP-hphMX6 ura4-294* | Microscopy, Rec25 localization, plasmid-borne *rec10* |
| GP9836 | *h*$^{90}$ *rec10-301::GFP-natMX6 ura4-D18* | Used to create GP9845 |
| GP9845 | *h*$^{90}$ *rec10-306::ura4*$^{+}$*-GFP-natMX6 ura4-D18* | Used to create GP9876 – GP9889 |
| GP9852 | *h*$^{+}$ *ade6-M26 ura4-D18 arg1-14 rec10-301::GFP-natMX6* | Recombination assay, chromosomal *rec10::GFP* |
| GP9859 | *h*$^{-}$ *rec10-301::GFP-natMX6 pat1-as1*(L95G)*::kanMX6* | Western blot, Rec10 abundance; |
| GP9860 | *h*$^{-}$ *pat1-as1*(L95G)*::kanMX6* | Western blot, Rec10 abundance; |
| GP8762 | *h*$^{90}$ *rec10-203::GFP-kanMX6* | Used to create GP9836 |
| GP7301 | *h*$^{-}$ *ade6-52 ura4-D18 rec10-260::ura4*$^{+}$ | Used to create GP9989 – GP9995; microscopy, Rec25 localization |
| GP8766 | *h*$^{90}$ *rec25-204::GFP-kanMX6* | Used to create GP9745 |
| GP9745 | *h*$^{90}$ *rec25-303::GFP-hphMX6* | Microscopy, Rec25 localization, chromosomal *rec10* |

[a] Additional strains are in Table S4.

[b] Strains were constructed by standard matings (Smith, 2009) or as described in Methods. Genealogies are available upon request. Sources of alleles, other than commonly used auxotrophies and *mat*, are the following: *pat1-as1*(L95G)*::kanMX6* (Guerra-Moreno et al., 2012); *rec10-175::kanMX6* (Ellermeier and Smith, 2005); *rec10-203::GFP-kanMX6* (Fowler et al., 2013); *rec10-260::ura4*$^{+}$ (Ma et al., 2017) *rec10-301::GFP-natMX6* (Materials and Methods); *rec10-306::ura4*$^{+}$*-GFP-natMX6* (Materials and Methods); *rec25-204::GFP-kanMX6* (Davis et al., 2008); *rec25-303::GFP-hphMX6* (Materials and Methods).

**Table S2. Plasmids**[a]

| Plasmid | Genotype | Ref. or origin |
|---|---|---|
| pFY20 | *ura4*$^{+}$ *ars1 stb amp* | Li et al. (1997) |
| pYL176 | *rec10*$^{+}$ *URA3*[b] *ars1 amp* | Lin and Smith (1995) |
| pFA6a-*natMX6* | *natMX6* | Hentges et al. (2005) |
| pFA6a-*hphMX6* | *hphMX6* | Hentges et al. (2005) |

[a] Additional plasmids are listed in Table S4.

[b] *S. cerevisiae URA3,* which complements *S. pombe ura4* mutations.

**Table S3. Oligonucleotides for *rec10* NLS mutant constructions**

| Oligo number | Nucleotide sequence (5' → 3') |
|---|---|
| OL4134 | AGATGGAAAGTTTGCAAAATCGACACAAAAATCTTTAAAACCTGATACTG |
| OL4135 | CAGTATCAGGTTTTAAAGATTTTTGTGTCGATTTTGCAAACTTTCCATCT |
| OL4136 | CTGAAAATCAAGAATCTTCGGTGGCGAAATCCAATGTTAATTTGCA |
| OL4137 | TGCAAATTAACATTGGATTTCGCCACCGAAGATTCTTGATTTTCAG |
| OL4140 | GCTGCACAAAAATCTTTAAAACCTGATAC |
| OL4141 | TGCAGCTGTCGATTTTGCAAACTTTC |
| OL4142 | GCTGCAGCGAAATCCAATGTTAATTTG |
| OL4143 | TGCAGCCACCGAAGATTCTTGATTTTC |
| OL4154 | CACTTCCAAGCAAGCATCCCAG (used for PCR amplification of NLS region) |
| OL4147 | CCTGTACTCAAGTTCCTGGCGA (used for PCR amplification of NLS region) |
| OL4155 | GACAAGAGTGTGTGCGACGATG (used for sequencing PCR products) |
| OL1780 | GTAACCGTCACTTATCGATGG (used for PCR analysis of *rec10*$^+$ and *rec10-306*) |
| OL1781 | AGCATGGACAGTATTGGCAAC (used for PCR analysis of *rec10*$^+$) |
| OL2124 | ATGCTCCTACAACATTACCAC (used for PCR analysis of *rec10-306*) |
| OL4278 | CACGCACAATCAACTGAAAC (*rec10* left forward primer to make *rec10-306*) |
| OL4279 | TTTCGTCAATATCACAAGCTCGGCAGTTCAATTTCTTGC (*rec10* left *ura4* flank reverse primer to make *rec10-306*) |
| OL4280 | GTGGGATTTGTAGCTAAGCTCCTACGATAGCAAACATTGC (*rec10* right *ura4* flank forward primer to make *rec10-306*) |
| OL4281 | TCCTGTACTCAAGTTCCTGG (*rec10* right reverse primer to make *rec10-306*) |
| OL4282 | GCAAGAAATTGAACTGCCGAGCTTGTGATATTGACGAAA (*rec10::ura4*$^+$ forward primer to make *rec10-306*) |
| OL4283 | GCAATGTTTGCTATCGTAGGAGCTTAGCTACAAATCCCAC (*rec10::ura4*$^+$ reverse primer to make *rec10-306*) |

**Table S4. *rec10* nuclear localization signal (NLS) mutants**

| *rec10* allele | Alternate designation | Amino acid sequence[a] Site A … Site B | Oligos; plasmid recipient | Plasmid isolate | Chromosomal isolate[b] | Chromosomal isolate[c] |
|---|---|---|---|---|---|---|
| *301* | + | **KRKK**QKSLKPDTENQESSV**KNKK** | | | GP9836 | |
| *289* | ΔA | ΔΔΔΔ------------------- | OL4134, OL4135 pYL176 | pMW2 | GP9876 | GP9992 |
| *290* | ΔB | -------------------ΔΔΔΔ | OL4136, OL4137 pYL176 | pMW3 | GP9877 | GP9989 |
| *296* | ΔA ΔB | ΔΔΔΔ---------------ΔΔΔΔ | OL4134, OL4135 pMW3 | pMW9 | GP9886 | GP9993 |
| *292* | AlaA | AAAA------------------- | OL4140, OL4141 pYL176 | pMW5 | GP9878 | GP9990 |
| *293* | AlaB | -------------------AAAA | OL4142, OL4143 pYL176 | pMW6 | GP9879 | GP9995 |
| *299* | AlaA AlaB | AAAA---------------AAAA | OL4140, OL4141 pMW6 | pMW12 | GP9889 | GP9991 |

[a] Rec10 amino acids from 497 – 519. The wild-type sequence is given for the + allele (*rec10*$^+$ on the plasmid and *rec10*$^+$ or *rec10-301::GFP-natMX6* on the chromosome). For other alleles "-" indicates the amino acid is that of wild type, and "Δ" or "A" indicates the amino acid is deleted or changed to Ala, respectively.

[b] Strains with the *rec10-NLS-GFP* fusion alleles. Strains other than GP9836 are derived from 5-fluoro-orotic acid-resistant (FOA$^R$) transformants of strain GP9845. See Materials and methods.

[c] Strains with the *rec10-NLS* alleles without the GFP fusion. Strains are derived from FOA$^R$ transformants of strain GP7301. See Materials and methods.

### Supplemental references


Davis, L., Rozalén, A.E., Moreno, S., Smith, G.R., and Martin-Castellanos, C. (2008). Rec25 and Rec27, novel components of meiotic linear elements, link cohesin to DNA breakage and recombination in fission yeast. Current Biology *18*, 849-854.

Ellermeier, C., and Smith, G.R. (2005). Cohesins are required for meiotic DNA breakage and recombination in *Schizosaccharomyces pombe*. Proc. Natl. Acad. Sci. USA *102*, 10952-10957.

Fowler, K.R., Gutiérrez-Velasco, S., Martín-Castellanos, C., and Smith, G.R. (2013). Protein determinants of meiotic DNA break hotspots. Molecular Cell *49*, 983-996.

Guerra-Moreno, A., Alves-Rodrigues, I., Hidalgo, E., and Ayte, J. (2012). Chemical genetic induction of meiosis in *Schizosaccharomyces pombe*. Cell Cycle *11*, 1621-1625.

Hentges, P., Van Driessche, B., Tafforeau, L., Vandenhaute, J., and Carr, A.M. (2005). Three novel antibiotic marker cassettes for gene disruption and marker switching in *Schizosaccharomyces pombe*. Yeast *22*, 1013-1019.

Li, Y.F., Numata, M., Wahls, W.P., and Smith, G.R. (1997). Region-specific meiotic recombination in *S. pombe*: the *rec11* gene. Molecular Microbiology *23*, 869-878.

Lin, Y., and Smith, G.R. (1995). Molecular cloning of the meiosis-induced *rec10* gene of *Schizosaccharomyces pombe*. Current Genetics *27*, 440-446.

Ma, L., Fowler, K.R., Martin-Castellanos, C., and Smith, G.R. (2017). Functional organization of protein determinants of meiotic DNA break hotspots. Scientific Reports *7*, 1393.

Smith, G.R. (2009). Genetic analysis of meiotic recombination in *Schizosaccharomyces pombe*. In Meiosis, S. Keeney, ed. (Totowa, NJ: Humana Press), pp. 65-76.
